# The genetic basis of extra-lobed leaves in ivyleaf morning glory (*Ipomoea hederacea*)

**DOI:** 10.64898/2026.10.27.762295

**Authors:** Martin R. Henry, Damian J. Hernandez, Margaret C. Li, Emily Glasgow, Amanda L. Peake, John R. Stinchcombe

**Affiliations:** Department of Ecology and Evolutionary Biology, University of Toronto 25 Willcocks Street, Room 3055, Toronto, Ontario, Canada, M5S 3B2

**Keywords:** leaf shape, bulked segregant analysis, leaf lobing, QTLseqr, QTL

## Abstract

Intraspecific leaf shape variation is common among angiosperms. Ivyleaf morning glory (*Ipomoea hederacea*) has become a useful system for studying the genetics and ecology of leaf shape variation as this species exhibits two leaf shapes: leaves with entire margins, and 3-lobed leaves. It is known that these phenotypes are controlled by a single locus, and differ in their ecophysiology, geographic distribution, ecology, and fitness. However, while lobed individuals producing leaves with 1-2 extra lobes have also been regularly observed, the genetic basis, if any, of these phenotypes has not been described. To investigate the genetic basis of extra-lobing, we compared phenotypic distributions of leaf shapes of an F2 generation to Mendelian ratios for a single locus trait and conducted a bulked segregant analysis to uncover quantitative trait loci. Our results suggest that producing extra-lobed leaves has a complex genetic basis and that there are multiple QTLs underlying this phenotype containing potential candidate genes involved with auxin distribution and leaf development. Whether these extra-lobed phenotypes also differ in terms of their physiology and ecology remains unexplored.

**Article summary:** Ivyleaf morning glory exhibits two main leaf shapes, heart-shaped entire leaves and 3-lobed, controlled by a single Mendelian locus. However, within the lobed genotypes, there are occasional individuals that produce leaves with extra lobes. We examined whether there was a simple genetic basis to producing extra lobes, and if so, does it have the same genetic basis as the other leaf shapes, or are different regions in the genome responsible? After planting 500 individuals in a greenhouse and collecting phenotypic and genomic data, we found that extra-lobing is heritable, does not have a simple Mendelian basis, but may be influenced by multiple QTLs.

## Introduction

Leaves display tremendous diversity across angiosperms. Leaves broadly fall into two general categories: simple (e.g., *Arabidopsis thaliana*) or compound (having multiple small leaflets per leaf; e.g., *Solanum lycopersicum*). Leaves can also be further described as having entire (smooth) or non-entire (e.g. serrated, lobed, etc.) margins. Variation in leaf morphology including leaf dissection, marginal structures, size, thickness, venation, and trichome density influence how plants photosynthesize, intercept light, regulate heat, support the leaf structure, use water efficiently, and avoid herbivory (Brown et al. 1991; Givnish 1979; Nicotra et al. 2011; Vogel 1970). Here, we investigate the genetic basis of a previously uncharacterized leaf shape phenotype in ivyleaf morning glory (*Ipomoea hederacea*), identifying multiple QTLs associated with the production of four and five lobed leaves, including regions that contain candidate genes involved with auxin distribution and leaf development.

Intraspecific leaf shape variation is common, and it is often assumed that differences in leaf shape are ecologically important adaptations to differences in environmental conditions across a species range (Givnish 1987). For example, leaf shape in butterfly weed (*Asclepias tuberosa*) differs between eastern and western populations, and this difference was correlated with moisture stress (Wyatt and Antonovics 1981). Red maple (*Acer rubrum*) leaves were found to be more dissected and toothed at colder locations, and California black oak (*Quercus kelloggii*) leaves were also more toothed with smaller leaf areas at higher elevations (Royer et al. 2008). In addition, *Mimulus laciniatus* was also shown to have more deeply lobed leaves at higher elevations (Love and Ferris 2024). Some studies have found that leaf shape appears to be under selection, but few have demonstrated a clear selective agent (Bright and Rausher 2008; Campitelli and Stinchcombe 2013; Ferris and Willis 2018; Nagy 1997; Richards et al. 2019). One challenge is that leaf shape may be correlated with other functional traits that are the actual targets of selection (Nagy 1997). Identifying the genetic basis of leaf shape variation may help clarify whether selection is acting directly on the loci responsible for shape itself, correlated pleiotropic effects of anatomical or physiological consequences of leaf shape, or traits that are genetically linked to leaf shape loci (Peake et al. 2026).

Leaf shape is the outcome of a complex interaction between hormonal physiology, genetics, and environmental factors. Leaves begin as leaf primordia on the shoot apical meristem where high auxin concentrations are generated by PIN-FORMED (PIN) auxin efflux carrier proteins (Benková et al. 2003; Heisler et al. 2005; Petrášek et al. 2006; Scarpella et al. 2010). The leaf will then undergo primary morphogenesis, beginning with the critical establishment of the adaxial-abaxial, or upper and lower, leaf identities. Marginal structures such as serrations and lobes develop early on in areas of high auxin concentration driven by PIN1-dependant polar auxin transport, in conjunction with CUP-SHAPED COTYLEDON (CUC) transcription factors and microRNA (miRNA) coordination in the leaf sinuses (Benková et al. 2003; Bilsborough et al. 2011; Nikovics et al. 2006). In secondary morphogenesis the leaf has its basic shape, but its ultimate form will come from cellular expansion, and the development of mature vasculature and epidermal cell types such as guard cells and trichomes (Efroni et al. 2010; Kalve et al. 2014). The development of more complex leaves (i.e. more leaflets, deeper lobes, etc.) is in part due to prolonged cell proliferation and expansion in localized areas on the leaf margin (Barth et al. 2009; Hagemann and Gleissberg 1996; Ori et al. 2007). Most angiosperm leaves are determinate structures and will eventually cease growth to achieve a final shape and size.

*Ipomoea hederacea* (L.) Jaquin (Convolvulaceae) shows a well-known Mendelian polymorphism for leaf shape (3-lobed versus heart-shaped, entire leaves; Figure 1) that appears to be under selection and follows a latitudinal cline (Bright and Rausher 2008; Bright-Emlen 1998; Campitelli and Stinchcombe 2013; Elmore 1986). However, it is not uncommon to observe lobed genotypes producing leaves with either 4 or 5 lobes (Figure 1). The phenotype is only observed in the lobed genetic background, suggesting either genotype by environment or epistatic interactions between the Mendelian leaf shape locus and unknown loci and/or environmental cues. While the genetics, ecology, and evolution of the 3-lobed polymorphism has been intensely investigated (Boyle et al. 2024; Bright and Rausher 2008; Bright-Emlen 1998; Campitelli et al. 2008, 2013; Campitelli and Stinchcombe 2013, 2014; Elmore 1986; Ennos 1981; Hull-Sanders et al. 2005; Peake et al. 2026; Singhal et al. 2025; Stinchcombe 2002; Stinchcombe et al. 2001), the ecological significance and the genetic basis of the extra-lobed phenotype remains unknown.

**Figure 1:**
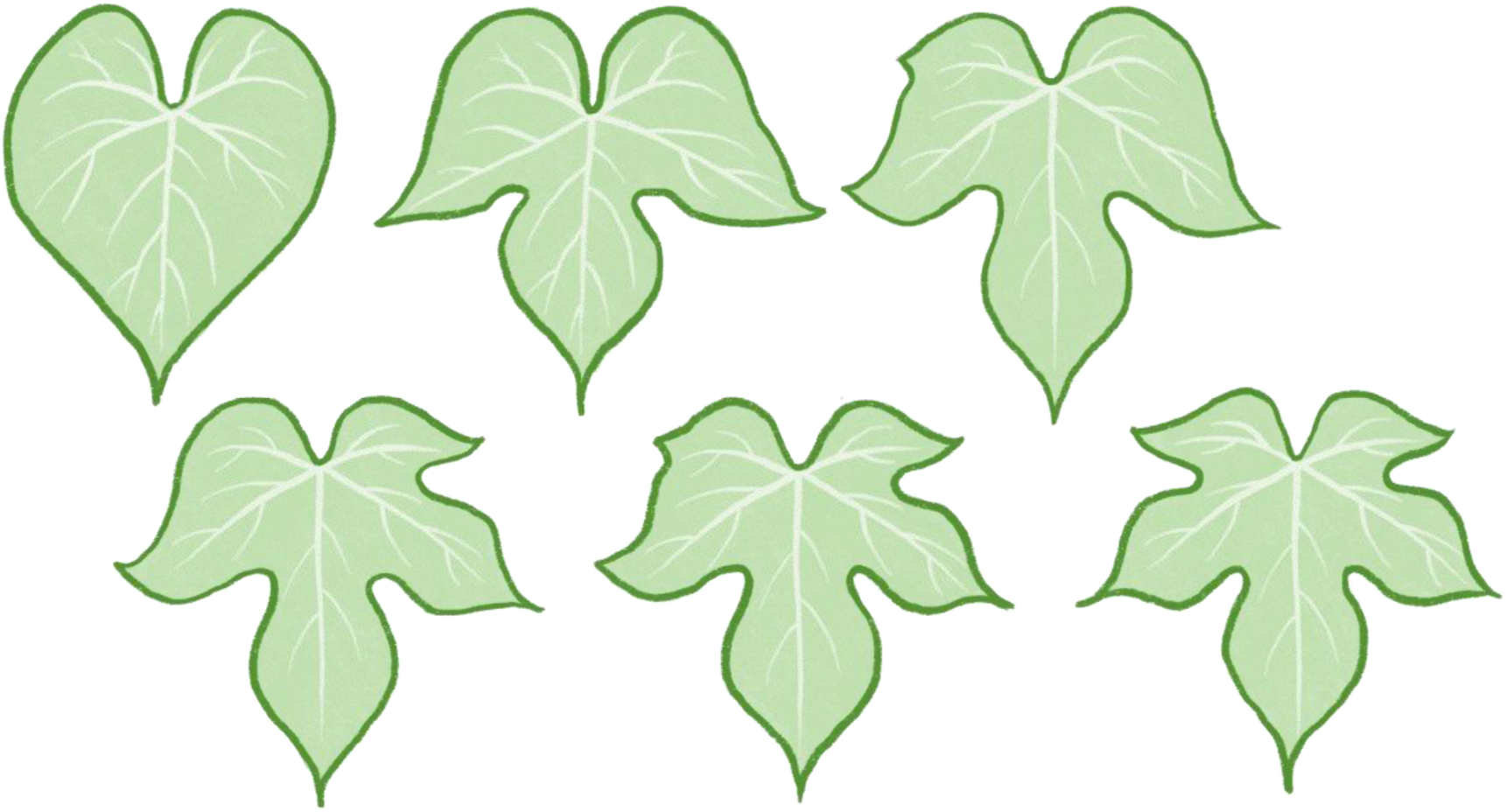
Leaf shape in *I. hederacea*: entire, 3-lobed, weakly 4-lobed, 4-lobed, weakly 5-lobed, fully 5-lobed (from left to right, top to bottom). Drawing by Martin Rose Henry.

In this study, we examined the genetic basis of extra-lobing in *I. hederacea* using a combination of controlled crosses and next-generation bulked segregant analysis (BSA). We identified quantitative trait loci (QTLs) that are associated with the extra-lobed phenotype using a BSA. Additionally, we mapped *I. hederacea* genes to *A. thaliana* and *S. lycopersicum* reference genome annotations to determine which genes and protein functions are present in significantly different QTL regions. We also conducted a gene ontology (GO) enrichment analysis to identify putative functions that are more strongly enriched in extra-lobed plants relative to 3-lobed plants. With the combination of gene identification through genome-wide protein alignments and GO enrichment, we identified potentially important genes underlying the extra-lobed phenotype in ivyleaf morning glory.

## Materials and Methods

### Species description

Ivyleaf morning glory, *I. hederacea*, is a weedy annual flowering vine found in eastern North America. It inhabits disturbed sites, roadside ditches, and crop fields (Bright-Emlen 1998; Mohr 1901; Shreve et al. 1910; Stevens 1948). Ivyleaf morning glory frequently self-pollinates, with selfing rates of up to 93% in some populations (Campitelli and Stinchcombe 2014; Ennos 1981).

### Crossing design, greenhouse methods, and phenotyping

We planted 500 F_2_ individuals derived from a cross between a 3-lobed individual and an individual regularly producing 5-lobed leaves. Parental lines had been collected from Delaware [39.721049 °N, - 75.745283 °W] and Maryland [38.83662 °N, -76.07692°W] respectively, and had been self-fertilized for 3 generations prior to the cross. We allowed a single F_1_ individual to self-fertilize to produce F_2_ experimental seeds. We then grew F_2_ plants in 26 cm^3^ tree pots and top watered them until they germinated. Benches were then flooded on a 6-hour schedule, with a 16-hour daylength and at 26-30°C°. Further details on growth conditions are in Supplementary Materials.

After one month, we scored the leaf shape phenotype for each plant over three days. We scored plants as: 3-lobed; 4-lobed if they had at least one leaf with an extra lobe; 4- and 5-lobed if they had at least one of each; and 5-lobed if they had at least one leaf with 5-lobes, but no 4-lobed leaves.

### DNA extraction and genomic sequencing

We sampled small, fresh leaves for DNA extraction from 142 of the 500 plants: 71 that were 3-lobed, and 71 that produced leaves with extra lobes (i.e., the presence of at least one 4- or 5-lobed leaf) as there was only a single individual in the 5-lobed category; we refer to the 4/5-lobed bulk as the extra-lobed bulk. We stored tissue at -80℃ prior to DNA extraction and used the Qiagen DNEasy Plant Mini Prep Kit for DNA extraction. We followed the manufacturer’s protocol except for the last step where we used 50µl instead of 100µl of Buffer AE for elution to get a more concentrated final product. Each pool contained equal concentrations of DNA of each individual sample. The Centre for Applied Genomics at SickKids Hospital performed library preparation and paired-end sequencing over two lanes (Illumina NovaSeqX 2x150bp paired-end sequencing).

### Genetic Analysis and Bioinformatics

#### Mendelian analysis

We first tested whether a simple 1- or 2-locus Mendelian model could explain the distribution of extra-lobed leaves. We used a chi-squared test to determine if the phenotypic distribution was significantly different from expectations for a simple Mendelian trait. Based on the presence of extra-lobing in the F_1_, we hypothesized that it was the dominant phenotype for a 3:1 expectation, and considered the 4-lobed phenotype as the putative heterozygous phenotype for testing 1:2:1 expectations. We considered testing a two-locus model and 9:3:3:1 expectation, with 3-lobed, 4-lobed, 4/5-lobed and only 5-lobed as the respective categories, but did not because we only observed a single individual in the last category.

#### Bioinformatics

For genomic data and BSA, unless otherwise indicated, all data was analyzed using high-performance computers from the Digital Research Alliance of Canada. We decompressed ORA files from SickKids using Illumina’s ORA decompression software (orad version 2.6.1, Illumina). To assess sequencing quality, we first analyzed the decompressed reads with FastQC (version 0.12.1)(Andrews 2010). Initial trimming was done using Cutadapt (version 5.1) within the Trim Galore wrapper (version 0.6.10), which also automatically ran FastQC for assessing read quality after trimming (Krueger et al. 2023; Martin 2011). Reads with a PHRED quality score lower than 20 were removed. We also removed the last 5 bp of all reads to account for an overrepresentation of poly-G tails. Summary statistics for sequencing depth are presented in Table 1.

**Table 1:** Summary of Illumina NovaSeqX sequencing data.

| **Sample** | **3-lobed** | **Extra-lobed** |
|---|---|---|
| Number of mapped reads | 320,399,224 (99.87%) | 364,285,564 (99.87%) |
| Mean depth | 66.0866x | 75.1754x |
| Coverage standard deviation | 275.6161 | 312.033x |
| Coverage >= 1x (%) | 99.4 | 99.46 |
| Coverage >= 5x (%) | 98.8 | 98.92 |
| Coverage >= 10x (%) | 98.25 | 98.45 |
| Coverage >= 40x (%) | 89.58 | 92.76 |

We aligned reads to the *I. hederacea* reference genome (Peake et al. 2026) using *bwa-mem2* (version 2.2.1)(Vasimuddin et al. 2019). We sorted, indexed, and marked duplicates in the resulting bam alignment files using *SAMtools* (version 1.20)(Danecek et al. 2021). We subsampled reads from the extra-lobed pool to match the sequencing depth of the 3-lobed pool, as discrepancies in read depth can affect downstream tests (Mansfeld and Grumet 2018). We then identified SNPs for each pool using the *HaplotypeCaller* function from the Genome Analysis Toolkit (*GATK*, version 4.6.1.0) and then combined the individual SNP files (vcf files) with the CombineGVCFs function (Van der Auwera et al. 2013). We converted this combined SNP matrix into a comma-separated table (*VariantsToTable* function) compatible with the *QTLseqR* pipeline used to perform the BSA in *R* (version 4.3.3, R Core Team 2024).

#### Bulked segregant analysis

We used *QTLseqR* (version 0.7.5.2) to conduct the BSA (Mansfeld and Grumet 2018) which implements both the QTL-seq (Takagi et al. 2013) and the G’ (Magwene et al. 2011) methods. QTL-seq calculates a Δ(SNP-index) over a 10 Mb sliding window (Takagi et al. 2013). The Δ(SNP-index) is the absolute difference in the ratio of reads that differ from the reference genome to the total number of reads at that position between the two bulks. Larger absolute differences in Δ(SNP-index) suggest genes or loci in that region influence the trait and differ between the two bulks, and should approach zero for unlinked loci that are not associated with the phenotype. The original Takagi method averages Δ(SNP-index) across a uniform window, but as suggested by the developers of *QTLseqR* we use a tricube-smoothed Δ(SNP-index), similar to the G’ method. We calculated 95% and 99% confidence intervals with *QTLseqr*, which randomly assigns alleles to two bulks in the absence of QTL and calculates Δ(SNP-index) to establish a null expectation (Mansfeld and Grumet 2018). We used 10,000 random samples to establish confidence intervals.

The G’ method calculates a G-statistic for every SNP (Magwene et al. 2011) G’ is then calculated by taking the tricube-weighted average of each SNP’s G-statistic within a 10Mb window to account for variation in sequencing coverage. A higher G’ value indicates greater allele frequency differences between the bulks. An FDR threshold of 0.01 was used as the significance threshold. We report the results for G’ in Supplementary Materials, and in the main text we present Δ(SNP-index) results with a focus on cases where the two methods identify the same QTL intervals.

### Gene mapping and GO enrichment analysis

We next identified candidate genes contributing to extra-lobing using the *I. hederacea* genome annotation (Peake et al. 2026) based on the results from the BSA. Loci with tricube-smoothed Δ(SNP-index) greater than the 95% confidence interval were selected for further investigation. We used the *map_snp_to_gene()* function from the *snpsettest* package (version 0.1.2) with a 20 kb window on either side of each gene (Joo and Himes 2023). We identified *I. hederacea* genes in the significant QTLs by finding the best-aligning protein-coding genes (lowest e-value, e-value cut-off of 0.001) within the *A. thaliana* genome (Araport 11, Cheng et al. 2017) and *S. lycopersicum* genome (ITAG 4.0, Hosmani et al. 2019). A full list of genes in QTL intervals is presented in Supplementary Table 1 with hypothesized candidate genes presented in the discussion.

Additionally, we identified functions significantly enriched in the QTLs-of-interest by conducting a GO enrichment analysis. Genes in *I. hederacea* inherited GO terms from their best gene detected in *A. thaliana* (i.e., the gene in *A. thaliana* with the lowest e-value). We conducted a GO enrichment analysis using a Fisher’s exact test (*topGO* package, version 2.54.0, Alexa and Rahnenfuhrer 2023) with the weight01 algorithm to account for the hierarchical nature of GO terms. The list of significant GO terms is in Supplemental Materials with those related to plant growth and development highlighted in the results.

## Results

### Phenotypes did not conform to simple Mendelian ratios

We observed extra-lobing in a common environment where F_2_ individuals had been randomized to their locations, suggesting that extra lobing can be observed even in the absence of G*E interactions. We next tested whether extra-lobing followed a simple Mendelian basis. Based on the observed phenotypic distributions, it is likely that extra-lobing in *I. hederacea* does not follow a simple 1- or 2-locus Mendelian model. Leaf-lobe phenotypes were almost uniformly distributed among offspring: 36.8% had only 3-lobed leaves, 34.6% produced at least some 4-lobed leaves, and 26.2% had 4-lobed leaves with at least one 5-lobed leaf (Table 2). The distribution of phenotypes differed significantly from the 1:2:1 expectation (ꭓ^2^ = 52.83, df = 2, P <0.0001) and 3:1 expectation (ꭓ^2^ = 42.01, df = 1, P <0.0001) (Figure 2) all significantly differed for both the 1:2:1 (χ^2^ = 52.83, df =2, P <0.0001) and 3:1 (χ^2^ = 42.01, df=1, P <0.0001) expectations.

**Figure 2:**
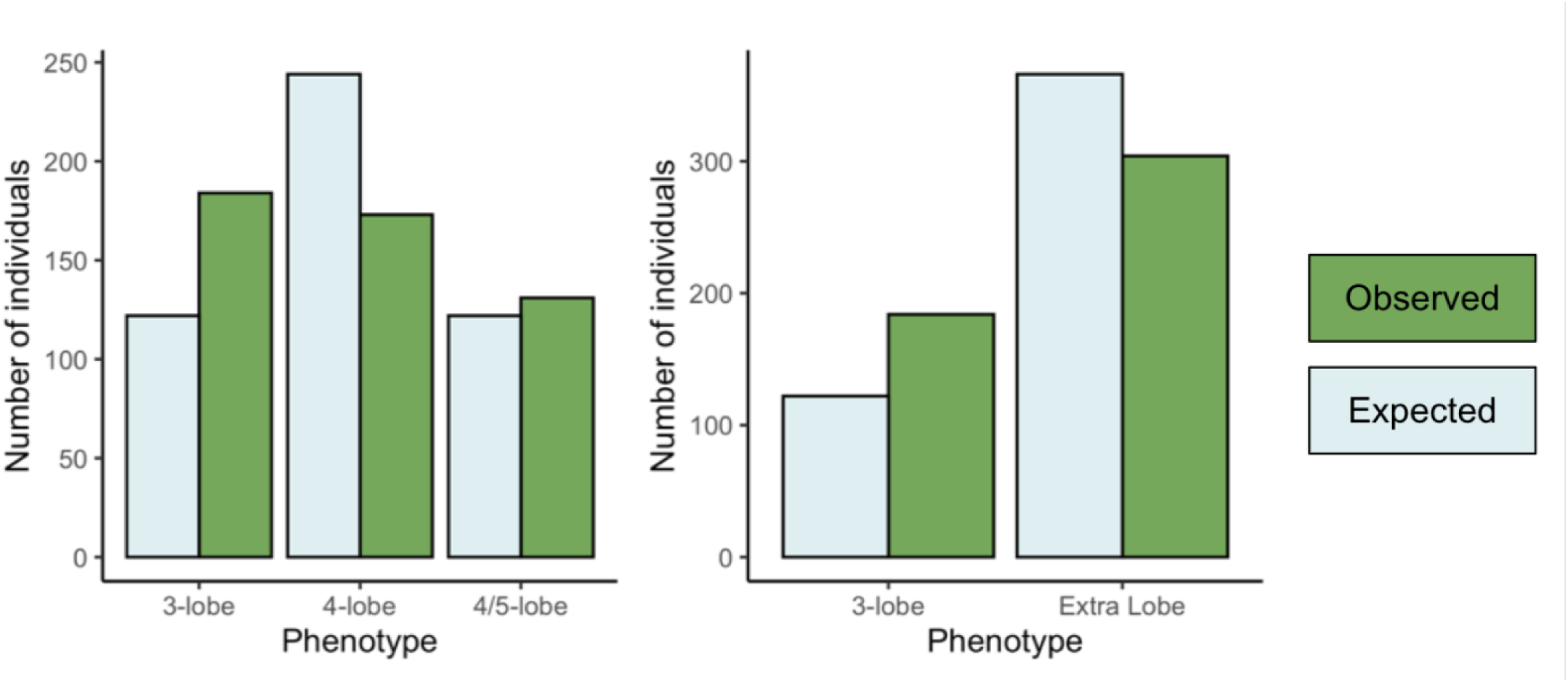
Phenotypic distributions for 1:2:1 and 3:1 Mendelian expectations. Green bars indicate observed frequencies, blue bars indicate expected frequencies. Chi-square tests between the observed and expected values

**Table 2:** Total leaf-shape phenotypes. Plants were scored as either 3-lobed, 4-lobed if they had at least one leaf with an extra lobe, 4- and 5-lobed if they had at least one of each, and 5-lobed if they had at least one leaf with 5-lobes.

| **Phenotype:** | **3-lobed** | **4-lobed** | **4- and 5- lobed** | **5-lobed** |
|---|---|---|---|---|
| Number of plants | 184 | 173 | 131 | 1 |
| Percentage | 36.8% | 34.6% | 26.2% | 0.2% |

### BSA identified loci underlying extra-lobed phenotype on two chromosomes

We found that there were significant differences between the 3-lobed and extra-lobed bulks associated with leaf shape, suggesting that the propensity for producing extra-lobing is heritable and that QTLs differing between the parental lines may underlie the phenotype. We identified two QTLs that show enrichment for SNPs over-represented in the extra-lobed bulk: one on Chromosome 2 and the other on Chromosome 4 (Table 3, Figure 3, Supplementary Figure 1, Supplementary Figure 2). Peake et al. (2026) recently identified a 117 kb insertion-deletion (indel) polymorphism on Chromosome 2 that perfectly co-segregates with entire and 3-lobed genotypes, where entire genotypes lack the 117 kb region. The QTL we detected, in a cross between between lobed genotypes which have the indel region intact, overlaps ∼ 52% of the indel region identified by (Peake et al. 2026) (Figure 4).

**Figure 3:**
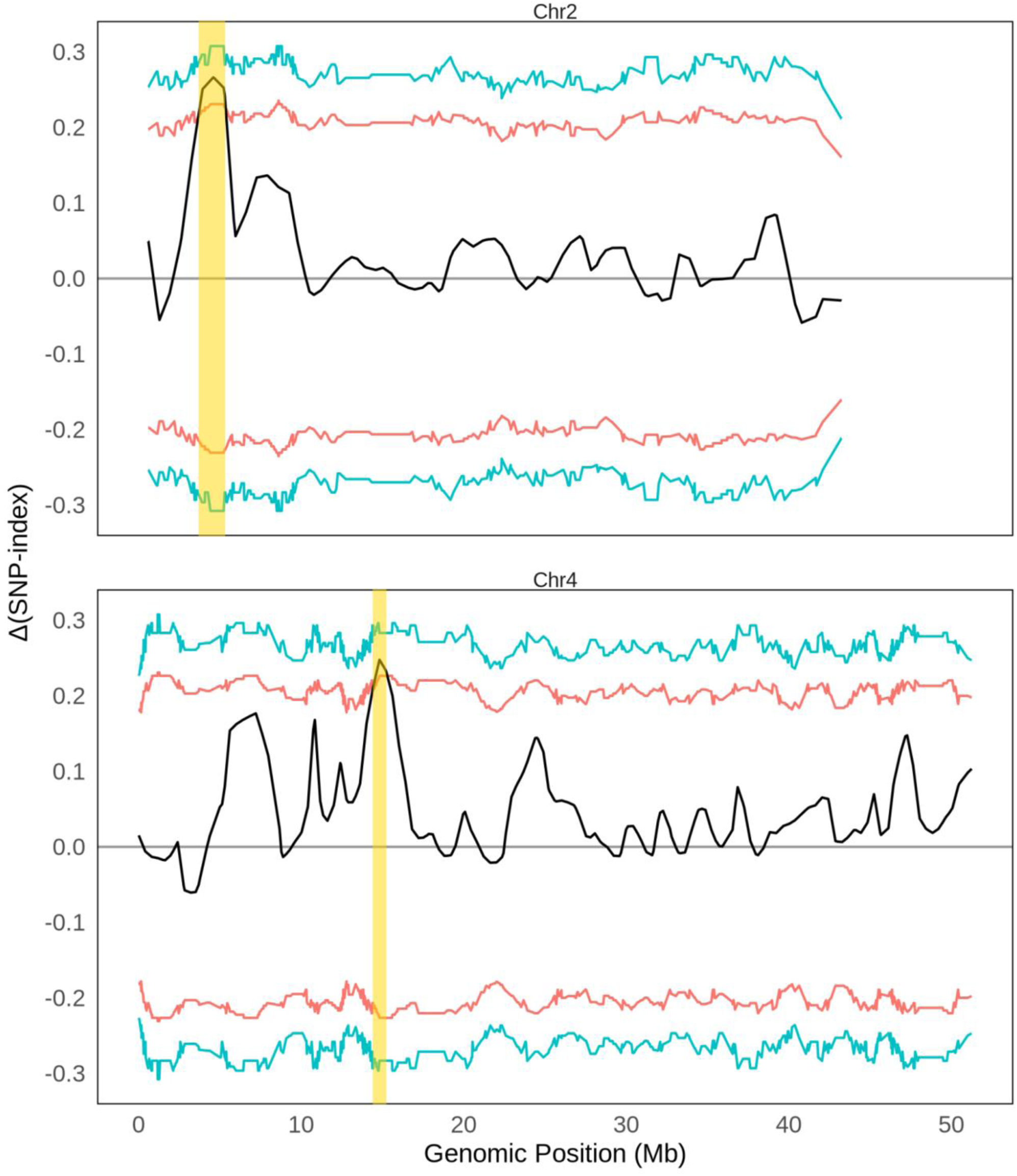
Plots showing Δ(SNP indices) across Chromosomes 2 and 4. Red line indicates a 95% confidence interval, blue line indicates 99% confidence interval. Yellow bar indicates significant regions.

**Figure 4:**
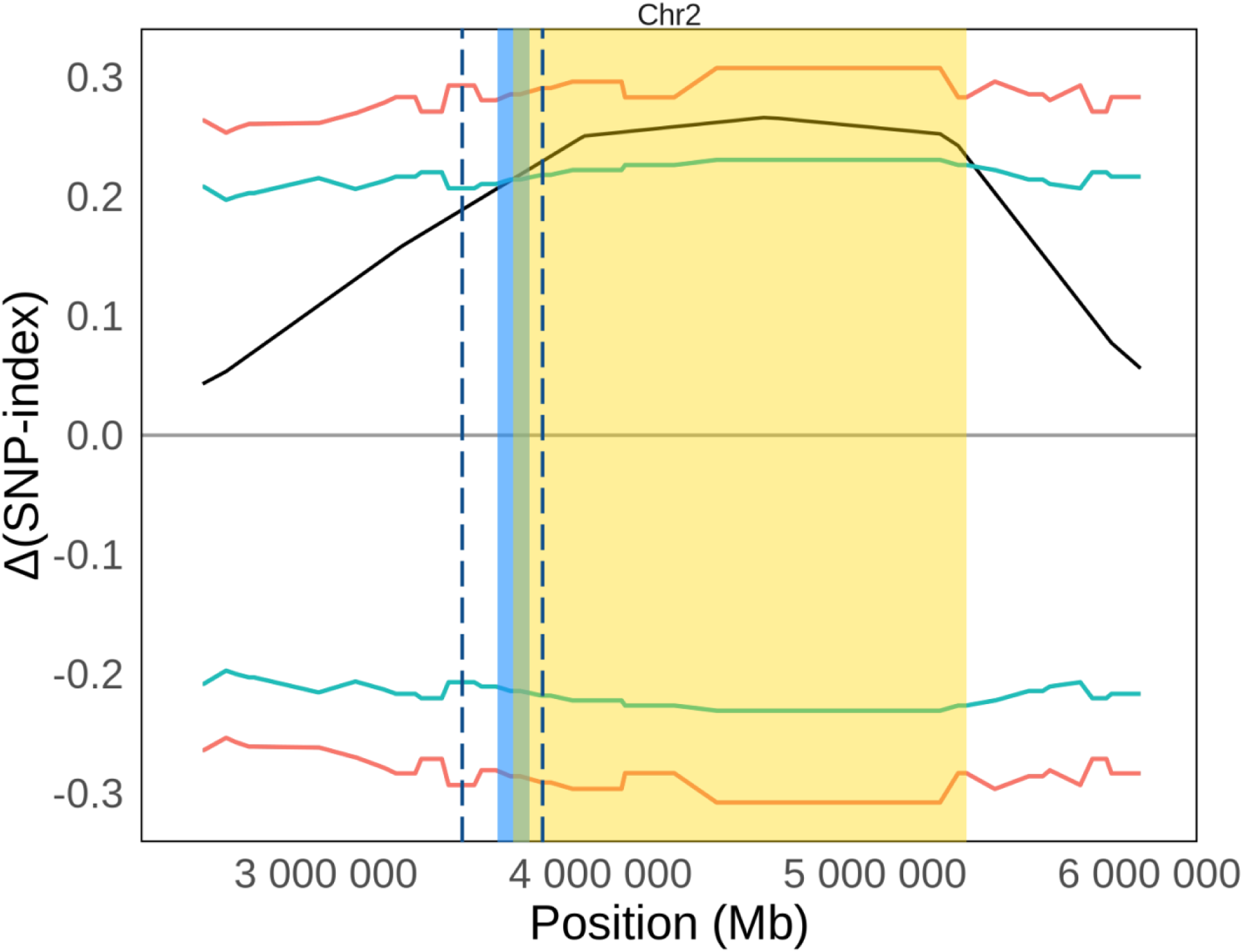
Zoom in on Chromosome 2, showing the QTL region identified in the BSA (yellow) next to the most significant SNPS in a GWAS for 3-lobed versus entire leaves (dotted lines) and the coordinates for the indel (light blue region), identified by (Peake et al. 2026). The green shaded interval indicates the overlap between the indel identified by Peake et al. and the QTL region identified here. Red line indicates a 95% confidence interval, blue line indicates 99% confidence interval.

**Table 3:**
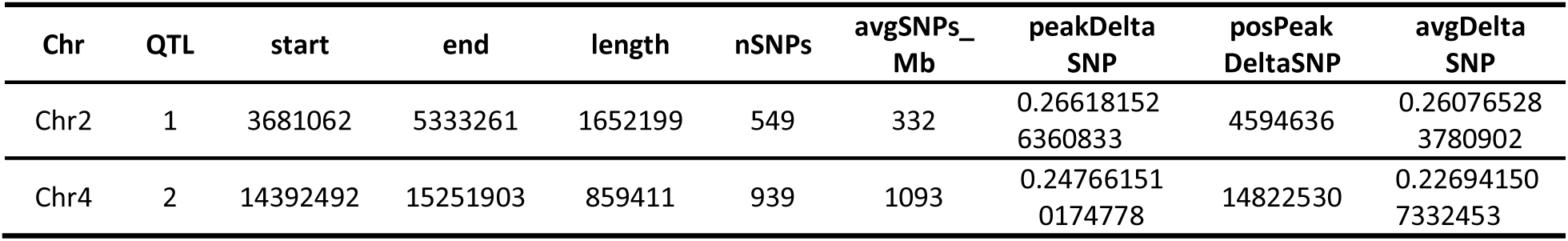
QTLs identified from bulked segregant analysis using the Δ(SNP index) calculated from the QTLseq method. CI = 95%.

| **Chr** | **QTL** | **start** | **end** | **length** | **nSNPs** | **avgSNPs_Mb** | **peakDeltaSNP** | **posPeakDeltaSNP** | **avgDeltaSNP** |
|---|---|---|---|---|---|---|---|---|---|
| Chr2 | 1 | 3681062 | 5333261 | 1652199 | 549 | 332 | 0.266181526360833 | 4594636 | 0.260765283780902 |
| Chr4 | 2 | 14392492 | 15251903 | 859411 | 939 | 1093 | 0.247661510174778 | 14822530 | 0.226941507332453 |

### QTLs contain genes involved in leaf growth and development

We found 98 gene IDs in the QTLs (Table 4, Supplementary Table 1). On Chromosome 2, we detected multiple genes involved in auxin distribution and response, as well as genes related to leaf polarity, cell expansion, sucrose synthase, and epidermal development, some of which also indirectly relate to auxin distribution and proper PIN protein expression (Table 5). These genes are *PIN-LIKES 1 (PILS1), AUXIN RESPONSE FACTOR 8 (ARF8), KANADI 1 (KAN1), EPIDERMAL PATTERNING FACTOR-LIKE 1 (EPFL1), EXPANSIN 13 (EXPA13), NAC28* [named after ***N****O APICAL MERISTEM* (*NAM*), ***A****RABIDOPSIS TRANSCRIPTION ACTIVATOR FACTOR* 1/2 (*ATAF1/2*), and ***C****UP-SHAPED COTYLEDON 2* (*CUC2*)], multiple *SUCROSE SYNTHASE (SUS)* genes*, HOMEODOMAIN GLABROUS 11 (HDG11),* and *ZWICHEL* (*ZWI)*. The QTL on Chromosome 4 contained a gene involved in regulating organ size, *BIG BROTHER (BB)*, and another involved in activating a KNOX gene, *NO TRANSMITTING TRACT (NTT*) (Table 5). Only 5 SNPs fell within the coordinates of any genes in the QTLs, one of which was a potential gene of interest: *HDG11*. We detail the specifics of the leaf development-related genes of interest in the Discussion.

**Table 4:** Number of gene IDs in the QTLs on Chromosomes 2 and 4, as well as how many mapped to *Arabidopsis thaliana* and *Solanum lycopersicum* genes.

|  | **Number of *I. hederacea* gene IDs** | **Number of *A. thaliana* genes** | **Number of *S. lycopersium* genes** |
|---|---|---|---|
| Chr2 | 63 | 68 | 76 |
| Chr4 | 35 | 34 | 37 |
| Total | 98 | 102 | 113 |

**Table 5:** List of hypothesized candidate genes, due to their known roles in leaf growth and development.

| Ipomoea hederacea gene ID | Chr | Arabidopsis thaliana accession ID | Arabidopsis thaliana gene | Solanum lycopersicum accession ID | Solanum lycopersicum gene |
|---|---|---|---|---|---|
| g17376 | Chr2 | AT1G65910 | NAC 28 | Solyc02g081270 | NAC domain-containing protein 53 (AHRD V3.3 *** A0A2G2XFS7_CAPBA) |
| g17378 | Chr2 | AT3G03220 | EXPANSIN 13 | Solyc03g031840 | expansin precursor 3 |
| g17379 | Chr2 | AT1G73360 | HOMEODOMAIN GLABROUS 11 | Solyc11g011940 | homeobox-leucine zipper protein ROC8 (AHRD V3.3 *** A0A2I4EBE3_9ROSI) |
| g17384 | Chr2 | AT5G16560 | KANADI 1 (KAN1) | Solyc11g011770 | Homeodomain-like superfamily protein (AHRD V3.3 *-* A0A2U1PJR7_ARTAN) |
| g17385 | Chr2 | AT4G02280 | SUCROSE SYNTHASE 3 | Solyc02g081300 | Sucrose synthase (AHRD V3.3 *** K4BA00_SOLLC) |
| g17385 | Chr2 | AT4G02280 | SUCROSE SYNTHASE 3 | Solyc03g098290 | Sucrose synthase (AHRD V3.3 *** K4BJP4_SOLLC) |
| g17385 | Chr2 | AT4G02280 | SUCROSE SYNTHASE 3 | Solyc09g098590 | Sucrose synthase (AHRD V3.3 *** E0Z1D1_SOLLC) |
| g17385 | Chr2 | AT5G37180 | SUCROSE SYNTHASE 5 | Solyc09g098590 | Sucrose synthase (AHRD V3.3 *** E0Z1D1_SOLLC) |
| g17385 | Chr2 | AT5G37180 | SUCROSE SYNTHASE 5 | Solyc02g081300 | Sucrose synthase (AHRD V3.3 *** K4BA00_SOLLC) |
| g17385 | Chr2 | AT5G37180 | SUCROSE SYNTHASE 5 | Solyc03g098290 | Sucrose synthase (AHRD V3.3 *** K4BJP4_SOLLC) |
| g17385 | Chr2 | AT1G73370 | SUCROSE SYNTHASE 6 | Solyc03g098290 | Sucrose synthase (AHRD V3.3 *** K4BJP4_SOLLC) |
| g17385 | Chr2 | AT1G73370 | SUCROSE SYNTHASE 6 | Solyc02g081300 | Sucrose synthase (AHRD V3.3 *** K4BA00_SOLLC) |
| g17385 | Chr2 | AT1G73370 | SUCROSE SYNTHASE 6 | Solyc09g098590 | Sucrose synthase (AHRD V3.3 *** E0Z1D1_SOLLC) |
| g17412 | Chr2 | AT5G37020 | AUXIN RESPONSE FACTOR 8 | Solyc03g031970 | Auxin response factor (AHRD V3.3 *** E0A8P3_SOLLC) |
| g17412 | Chr2 | AT5G37020 | AUXIN RESPONSE FACTOR 8 | Solyc02g037530 | Auxin response factor 8B |
| g17413 | Chr2 | AT5G65930 | ZWICHEL | Solyc02g068340 | Kinesin-like protein (AHRD V3.3 *** O24147_TOBAC) |
| g17419 | Chr2 | AT1G20925 | PIN-LIKES 1 | Solyc04g082830 | Auxin efflux carrier (AHRD V3.3 *** A0A200PR79_9MAGN) |
| g17420 | Chr2 | AT1G20925 | PIN-LIKES 1 | Solyc02g037550 | Auxin efflux carrier (AHRD V3.3 *** A0A2U1MR93_ARTAN) |
| g17446 | Chr2 | AT5G10310 | EPIDERMAL PATTERNING FACTOR-LIKE 1 | Solyc11g009040 | EPIDERMAL PATTERNING FACTOR-like protein 1 (AHRD V3.3 *** A0A2G2WTE5_CAPBA),Pfam:PF17181 |
| g23362 | Chr4 | AT3G63530 | BIG BROTHER | Solyc01g087060 | RING/U-box superfamily protein (AHRD V3.3 *** A0A2U1NQN4_ARTAN) |
| g23364 | Chr4 | AT3G57670 | NO TRANSMITTING TRACT | Solyc01g087050 | Zinc finger protein (AHRD V3.3 *** A0A200QA40_9MAGN) |

### GO enrichment analysis

We found 77 significant biological process GO terms and 40 significant molecular function GO terms. We report all of these in Supplementary Table 2 and Supplementary Table 3 for completeness. Here in the main text, we report the top 7 biological process and top 3 molecular function GO terms, which have the strongest P values (P < 0.00009) and approximately the top 9% and 7.5% of GO terms respectively, in addition to the GO terms described above with leaf shape candidate genes.

The top 7 biological process GO terms are related to nuclear-transcribed mRNA catabolic process, nonsense-mediated decay (GO:0000184), long-day photoperiodism (GO:0048571), RNAi-mediated antiviral immune response (GO:0009616), pollen-pistil interaction (GO:0009875), priming of cellular response to stress (GO:0080136), positive regulation of defense response (GO:1900426), and RNA splicing (GO:0008380). The top 3 molecular function GO terms are 5’-3’ DNA helicase activity (GO:0043139), RNA helicase activity (GO:0003724), and myosin binding (GO:0017022). In all of these cases, it is difficult to discern a relationship between the GO terms and leaf shape and morphology.

## Discussion

We identified multiple QTLs associated with extra-lobing in ivyleaf morning glory, suggesting that there is a genetic influence to the propensity of producing extra lobed leaves. Establishing that there is a genetic basis underlying the propensity to produce extra-lobed leaves is a necessary first step prior to future studies of its microevolutionary or ecological significance (Conner and Hartl 2004). In contrast to the well-studied entire -vs-3-lobed polymorphism in this species, the propensity for producing extra-lobed leaves did not follow Mendelian segregation patterns. Our BSA results suggest that multiple loci may be involved in the propensity for producing extra lobes, including a QTL that maps to the same region as the entire -vs-3-lobed polymorphism, suggesting that gene(s) in this region underlie leaf shape, morphology, and development in *I. hederacea*. Below, we discuss in detail some of the candidate genes present in these QTL intervals and future directions.

### Chromosome 2 Candidate Genes

We found multiple genes in the QTL on Chromosome 2 that are involved in auxin transport and auxin-inducible responses. The first is *PIN-LIKES 1* (*PILS1*) gene (GO:0080162, p = 0.005). *PILS* encodes for proteins that are hypothesized to have similar functions to PIN proteins in that they facilitate auxin transport and cellular response to auxin (Barbez et al. 2012; Sauer and Kleine-Vehn 2019). A second auxin-related gene in this QTL interval is *AUXIN RESPONSE FACTOR 8* (*ARF8*). *ARFs* are transcription factors that regulate auxin response genes and are involved in many aspects of plant growth and development (Li et al. 2016). Many *ARFs* work redundantly and share structural similarity (Okushima et al. 2005). *ARF8* specifically promotes proximal-distal leaf growth, and also represses Class-1 *KNOX* genes in floral organs, causing abnormal petal and vasculature development, and somewhat uneven leaf development upon floral initiation (Tabata et al. 2010; Y. Xiong et al. 2021). In tomato, *SlARF8* was also found to play a role in determining leaf morphology (Wu et al. 2018). A third gene, *KANADI 1* (*KAN1*), was also present in the QTL, and it specifies the development of the abaxial leaf identity (GO:0010158, p = 0.002)(Emery et al. 2003; Eshed et al. 2004; Kerstetter et al. 2001; Pekker et al. 2005). *KANADI* genes also influence the distribution of auxin within the developing embryo, regulates PIN expression, and vascular tissue development (Burian et al. 2022; Emery et al. 2003; Eshed et al. 2004; Guan et al. 2017; Ilegems et al. 2010; Izhaki and Bowman 2007). ARF3 and ARF4 also work in cooperation with *KANADI* genes to specify abaxial cell types (Pekker et al. 2005). Failure of either adaxial or abaxial identities to develop results in a lack of proper lamina outgrowth. For example, ectopic expression of *KANADI* genes in leaf primordia restricts marginal, and therefore lobe, outgrowth (Eshed et al. 2004). In Japanese morning glory (*Ipomoea nil*), mutations in *KAN1*, along with potentially other modifiers, cause improper establishment of the abaxial identity in lateral organs, leading to curled leaves and separated petals (Iwasaki and Nitasaka 2006). Lastly, *EPIDERMAL PATTERNING FACTOR-LIKE 1* (*EPFL1)* is involved in regulating shoot development, stomatal patterning, shoot apical meristem size, leaf initiation, and organ elongation (Bessho-Uehara et al. 2016; Bundy et al. 2012; Hara et al. 2007; Kosentka et al. 2019; Shimada et al. 2011). EPFL1 and 2 were both found to be necessary for the formation of sharp auxin maxima in developing leaves and the shoot apical meristem, as *epfl1* mutants have shallower leaf serrations, and *epfl1 epfl2* double mutants have broader auxin maxima (Tameshige et al. 2025).

Collectively, *EPFL1*, *KAN1*, *ARF8*, and *PILS1* represent promising candidate genes for variation in leaf morphology.

There are four other genes of interest present within the QTL on Chromosome 2 that may be related to leaf growth; for these genes, they belong to gene families that have members that have been demonstrated to be involved in leaf growth, development, or shape. We include these genes as candidates because *I. hederacea* is a non-model organism, and the genetic basis of leaf shape is often lineage specific (Nicotra, 2011). The first is *EXPANSIN A13* (*EXPA13*) for *A. thaliana*, or expansin precursor 3 in tomato. Expansins (*EXP*) belong to a large gene family with diverse and tissue-specific roles but are known to be involved with cell wall loosening (Cosgrove 2024). They are involved in leaf growth, as induced local expression of expansin in leaves causes outgrowths to develop (Pien et al. 2001). Again, auxin plays a part here: auxin reduces the pH of plant cells, and this acidic environment is required for expansin activity to occur (Cosgrove 2024). The second is *NAC28* or *NAC53* transcription factor in *A. thaliana* and tomato, respectively. NAC is a very large family of transcription factors. They are involved in diverse processes including embryogenesis, cotyledon development, biotic and abiotic stress responses, and lateral organ development (H. Xiong et al. 2025). This family contains a well-known player in leaf lobing formation, *CUC2*, which promotes leaf serrations in simple-leaved species like *A. thaliana* or more leaf dissections in compound species like tomato (Berger et al. 2009; Blein et al. 2008; Hasson et al. 2011). *NAC028* was found to be upregulated in more complex leaves of the heteroblastic species *Gevuina avellana* (Ostria-Gallardo et al. 2016; Wen et al. 2022). *NAC53* also encodes *ALADIN*, a protein involved in proper cell division, stem elongation, and plant development (Best et al. 2021). Third, we detected multiple *SUCROSE SYNTHASE (SUS)* genes, which may be involved in leaf development and auxin signalling, as tomatoes with *SlSUS1*, *SlSUS3* and *SlSUS4* mutations displayed deformed leaves and elevated *PIN1* levels (Goren et al. 2017). Fourth, this QTL interval contained *HOMEODOMAIN GLABROUS 11* (*HDG11*) for *A. thaliana,* or HD-zip protein ROC8 in tomato, which belongs to the Class IV HD-Zip family. *HDG11* was the only leaf development-related gene containing a SNP. The Class IV HD-Zip family is involved with the differentiation of specialized epidermal cells such as guard cells, pavement cells, and trichomes (Schrick et al. 2023). In Chinese kale, mutants that overexpressed *HDG11* had higher *YUCCA*, *EXPANSIN*, and *PIN* expression (Zhu et al. 2016).

### Chromosome 4 Candidate Genes

The majority of the genes on the QTL on Chromosome 4 do not have obvious relationships with leaf shape. However, there are two of note. *BIG BROTHER* (*BB*) encodes a RING finger protein and is a negative regulator of plant organ size by restricting the duration of cellular proliferation (Disch et al. 2006). Mutants experience increased leaf growth and a delayed transition from cellular proliferation to differentiation, and overexpression leads to significantly smaller leaf area (Vanhaeren et al. 2017). *NO TRANSMITTING TRACT* (*NTT*) is a zinc finger transcription that activates a Class 1 *KNOTTED-LIKE HOMEOBOX* (*KNOX*) gene, *BREVIPEDICELLUS* (*BP/KNAT1*). *NTT* has been studied more so for its role on fruit development, but overexpression or ectopic expression also causes irregular leaf margins and shorter internodes, similarly to when *BP/KNAT1* is overexpressed (Chuck et al. 1996; Chung et al. 2013; Lincoln et al. 1994; Marsch-Martínez et al. 2014).

## Conclusions and Future Directions

Nearly a century ago, classical work in *Ipomoea nil* (L.) Roth, the sister species to *I. hederacea*, attempted to discern the genetic basis of mutant four or five lobed phenotypes, observed in contrast to the typical 3-lobed phenotype (Imai 1938). The distribution of phenotypes observed by Imai (1938) also failed to conform to patterns of Mendelian segregation, and he hypothesized that modifier or suppressor genes were involved in this phenotype. Our results from *I. hederacea* also fail to detect a simple genetic basis for this phenotype and our BSA results suggest that multiple chromosomal regions may underlie this propensity.

While our work suggests that this phenotype does have a genetic influence, the ecological and evolutionary consequences of this shape variation– if any– remain unclear. Whether extra-lobing– either its presence or the number of extra-lobed leaves produced– confers differences in ecophysiology, geographic distribution, ecology, and fitness like the 3-lobed and entire phenotypes (Boyle et al. 2024; Bright and Rausher 2008; Bright-Emlen 1998; Campitelli et al. 2008, 2013; Campitelli and Stinchcombe 2013, 2014; Elmore 1986; Ennos 1981; Singhal et al. 2025; Stinchcombe 2002; Stinchcombe et al. 2001) warrants future investigation.

## Acknowledgements

We thank members of the Stinchcombe Lab for discussion and encouragement, Jannice Friedman for advice on BSA, and Tom Gludovacz and Alice DesRoches for horticultural support. We thank Jacqueline Sztepanacz, Asher Cutter, and Megan Bontrager for comments on previous versions of the manuscript.

## Data Availability Statement

The raw sequence reads supporting the findings of this study have been deposited in the NCBI SRA under BioProject PRJNA1346405. These data will be publicly released upon publication. All code can be found on GitHub.

## References

1. Alexa, Adrian, and Jorg Rahnenfuhrer. 2023. topGO: Enrichment Analysis for Gene Ontology. 10.18129/B9.bioc.topGO.

2. Andrews, Simon. 2010. FastQC: A Quality Control Tool for High Throughput Sequence Data. Released. http://www.bioinformatics.babraham.ac.uk/projects/fastqc.

3. Barbez, Elke, Martin Kubeš, Jakub Rolčík, et al. 2012. “A Novel Putative Auxin Carrier Family Regulates Intracellular Auxin Homeostasis in Plants.” Nature 485 (7396): 119–22. 10.1038/nature11001.

4. Barth, Sina, Thomas Geier, Klaus Eimert, Bernard Watillon, Rajbir S. Sangwan, and Stefan Gleissberg. 2009. “KNOX Overexpression in Transgenic Kohleria (Gesneriaceae) Prolongs the Activity of Proximal Leaf Blastozones and Drastically Alters Segment Fate.” Planta 230 (6): 1081–91. 10.1007/s00425-009-0997-0.

5. Benková, Eva, Marta Michniewicz, Michael Sauer, et al. 2003. “Local, Efflux-Dependent Auxin Gradients as a Common Module for Plant Organ Formation.” Cell 115 (5): 591–602. 10.1016/S0092-8674(03)00924-3.

6. Berger, Yael, Smadar Harpaz-Saad, Arnon Brand, et al. 2009. “The NAC-Domain Transcription Factor GOBLET Specifies Leaflet Boundaries in Compound Tomato Leaves.” Development 136 (5): 823–32. 10.1242/dev.031625.

7. Bessho-Uehara, Kanako, Diane R. Wang, Tomoyuki Furuta, et al. 2016. “Loss of Function at RAE2, a Previously Unidentified EPFL, Is Required for Awnlessness in Cultivated Asian Rice.” Proceedings of the National Academy of Sciences 113 (32): 8969–74. 10.1073/pnas.1604849113.

8. Best, Norman B., Charles Addo-Quaye, Bong-Suk Kim, et al. 2021. “Mutation of the Nuclear Pore Complex Component, *Aladin1*, Disrupts Asymmetric Cell Division in *Zea Mays* (Maize).” G3 Genes|Genomes|Genetics 11 (7): jkab106. 10.1093/g3journal/jkab106.

9. Bilsborough, Gemma D., Adam Runions, Michalis Barkoulas, et al. 2011. “Model for the Regulation of Arabidopsis Thaliana Leaf Margin Development.” Proceedings of the National Academy of Sciences 108 (8): 3424–29. 10.1073/pnas.1015162108.

10. Blein, Thomas, Amada Pulido, Aurélie Vialette-Guiraud, et al. 2008. “A Conserved Molecular Framework for Compound Leaf Development.” Science 322 (5909): 1835–39. 10.1126/science.1166168.

11. Boyle, Julia A., Megan E. Frederickson, and John R. Stinchcombe. 2024. “Genetic Architecture of Heritable Leaf Microbes.” Microbiology Spectrum 12 (7): e00610–24. 10.1128/spectrum.00610-24.

12. Bright, Kerry L., and Mark D. Rausher. 2008. “Natural Selection on a Leaf-Shape Polymorphism in the Ivyleaf Morning Glory (Ipomoea Hederacea).” Evolution 62 (8): 1978–90. 10.1111/j.1558-5646.2008.00416.x.

13. Bright-Emlen, Kerry Leigh. 1998. “Geographic Variation and Natural Selection on a Leaf Shape Polymorphism in the Ivyleaf Morning Glory (Ipomoea Hederacea).” Ph.D., Duke University. https://www.proquest.com/docview/304423201/abstract/1F38808984004F83PQ/1.

14. Brown, V. K., J. H. Lawton, and P. J. Grubb. 1991. “Herbivory and the Evolution of Leaf Size and Shape [and Discussion].” Philosophical Transactions: Biological Sciences 333 (1267): 265–72.

15. Bundy, Mark G. R., Olivia A. Thompson, Matthew T. Sieger, and Elena D. Shpak. 2012. “Patterns of Cell Division, Cell Differentiation and Cell Elongation in Epidermis and Cortex of Arabidopsis Pedicels in the Wild Type and in Erecta.” PLOS ONE 7 (9): e46262. 10.1371/journal.pone.0046262.

16. Burian, Agata, Gael Paszkiewicz, Khoa Thi Nguyen, Shreyas Meda, Magdalena Raczyńska- Szajgin, and Marja C. P. Timmermans. 2022. “Specification of Leaf Dorsiventrality via a Prepatterned Binary Readout of a Uniform Auxin Input.” Nature Plants 8 (3): 269–80. 10.1038/s41477-022-01111-3.

17. Campitelli, Brandon E., Amanda J. Gorton, Katherine L. Ostevik, and John R. Stinchcombe. 2013. “The Effect of Leaf Shape on the Thermoregulation and Frost Tolerance of an Annual Vine, Ipomoea Hederacea (Convolvulaceae).” American Journal of Botany 100 (11): 2175–82. 10.3732/ajb.1300136.

18. Campitelli, Brandon E., Anna K. Simonsen, Andrea Rico Wolf, Jessamyn S. Manson, and John R. Stinchcombe. 2008. “Leaf Shape Variation and Herbivore Consumption and Performance: A Case Study with Ipomoea Hederacea and Three Generalists.” Arthropod- Plant Interactions 2 (1): 9–19. 10.1007/s11829-007-9028-x.

19. Campitelli, Brandon E., and John R. Stinchcombe. 2013. “Natural Selection Maintains a Single- Locus Leaf Shape Cline in Ivyleaf Morning Glory, Ipomoea Hederacea.” Molecular Ecology 22 (3): 552–64. 10.1111/mec.12057.

20. Campitelli, Brandon E., and John R. Stinchcombe. 2014. “Population Dynamics and Evolutionary History of the Weedy Vine Ipomoea Hederacea in North America.” G3 Genes|Genomes|Genetics 4 (8): 1407–16. 10.1534/g3.114.011700.

21. Cheng, Chia-Yi, Vivek Krishnakumar, Agnes P. Chan, Françoise Thibaud-Nissen, Seth Schobel, and Christopher D. Town. 2017. “Araport11: A Complete Reannotation of the Arabidopsis Thaliana Reference Genome.” The Plant Journal 89 (4): 789–804. 10.1111/tpj.13415.

22. Chuck, G., C. Lincoln, and S. Hake. 1996. “KNAT1 Induces Lobed Leaves with Ectopic Meristems When Overexpressed in Arabidopsis.” The Plant Cell 8 (8): 1277–89. 10.1105/tpc.8.8.1277.

23. Chung, Kyung Sook, Jeong Hwan Lee, Jong Seob Lee, and Ji Hoon Ahn. 2013. “Fruit Indehiscence Caused by Enhanced Expression of NO TRANSMITTING TRACT in Arabidopsis Thaliana.” Molecules and Cells 35 (6): 519–25. 10.1007/s10059-013-0030-0.

24. Conner, Jeffrey K., and Daniel L. Hartl. 2004. A Primer of Ecological Genetics. Vol. 425. Sinauer Associates Sunderland, MA.

25. Cosgrove, Daniel J. 2024. “Plant Cell Wall Loosening by Expansins.” Annual Review of Cell and Developmental Biology 40 (1): 329–52. 10.1146/annurev-cellbio-111822-115334.

26. Danecek, Petr, James K. Bonfield, Jennifer Liddle, et al. 2021. “Twelve Years of SAMtools and BCFtools.” GigaScience 10 (2). 10.1093/gigascience/giab008.

27. Disch, Sabine, Elena Anastasiou, Vijay K. Sharma, Thomas Laux, Jennifer C. Fletcher, and Michael Lenhard. 2006. “The E3 Ubiquitin Ligase BIG BROTHER Controls Arabidopsis Organ Size in a Dosage-Dependent Manner.” Current Biology 16 (3): 272–79. 10.1016/j.cub.2005.12.026.

28. Efroni, Idan, Yuval Eshed, and Eliezer Lifschitz. 2010. “Morphogenesis of Simple and Compound Leaves: A Critical Review.” The Plant Cell 22 (4): 1019–32. 10.1105/tpc.109.073601.

29. Elmore, C. Dennis. 1986. “Mode of Reproduction and Inheritance of Leaf Shape in Ipomoea Hederacea.” Weed Science 34 (3): 391–95.

30. Emery, John F., Sandra K. Floyd, John Alvarez, et al. 2003. “Radial Patterning of Arabidopsis Shoots by Class III HD-ZIP and KANADI Genes.” Current Biology 13 (20): 1768–74. 10.1016/j.cub.2003.09.035.

31. Ennos, R. A. 1981. “Quantitative Studies of the Mating System in Two Sympatric Species of Ipomoea (Convolvulaceae).” Genetica 57 (2): 93–98. 10.1007/BF00131233.

32. Eshed, Yuval, Anat Izhaki, Stuart F. Baum, Sandra K. Floyd, and John L. Bowman. 2004. “Asymmetric Leaf Development and Blade Expansion in Arabidopsisare Mediated by KANADI and YABBY Activities.” Development 131 (12): 2997–3006. 10.1242/dev.01186.

33. Ferris, Kathleen G., and John H. Willis. 2018. “Differential Adaptation to a Harsh Granite Outcrop Habitat between Sympatric Mimulus Species.” Evolution 72 (6): 1225–41. 10.1111/evo.13476.

34. Givnish, Thomas J. 1979. “On the Adaptive Significance of Leaf Form.” In Topics in Plant Population Biology, edited by Otto T. Solbrig, Subodh Jain, George B. Johnson, and Peter H. Raven. Macmillan Education UK. 10.1007/978-1-349-04627-0_17.

35. Givnish, Thomas J. 1987. “Comparative Studies of Leaf Form: Assessing the Relative Roles of Selective Pressures and Phylogenetic Constraints.” New Phytologist 106 (s1): 131–60. 10.1111/j.1469-8137.1987.tb04687.x.

36. Goren, Shlomo, Nitsan Lugassi, Ofer Stein, et al. 2017. “Suppression of Sucrose Synthase Affects Auxin Signaling and Leaf Morphology in Tomato.” PLOS ONE 12 (8): e0182334. 10.1371/journal.pone.0182334.

37. Guan, Chunmei, Binbin Wu, Ting Yu, et al. 2017. “Spatial Auxin Signaling Controls Leaf Flattening in Arabidopsis.” Current Biology 27 (19): 2940–2950.e4. 10.1016/j.cub.2017.08.042.

38. Hagemann, Wolfgang, and Stefan Gleissberg. 1996. “Organogenetic Capacity of Leaves: The Significance of Marginal Blastozones in Angiosperms.” Plant Systematics and Evolution 199 (3): 121–52. 10.1007/BF00984901.

39. Hara, Kenta, Ryoko Kajita, Keiko U. Torii, Dominique C. Bergmann, and Tatsuo Kakimoto. 2007. “The Secretory Peptide Gene EPF1 Enforces the Stomatal One-Cell-Spacing Rule.” Genes & Development 21 (14): 1720–25. 10.1101/gad.1550707.

40. Hasson, Alice, Anne Plessis, Thomas Blein, et al. 2011. “Evolution and Diverse Roles of the CUP-SHAPED COTYLEDON Genes in Arabidopsis Leaf Development.” The Plant Cell 23 (1): 54–68. 10.1105/tpc.110.081448.

41. Heisler, Marcus G., Carolyn Ohno, Pradeep Das, et al. 2005. “Patterns of Auxin Transport and Gene Expression during Primordium Development Revealed by Live Imaging of the Arabidopsis Inflorescence Meristem.” Current Biology 15 (21): 1899–911. 10.1016/j.cub.2005.09.052.

42. Hosmani, Prashant S., Mirella Flores-Gonzalez, Henri van de Geest, et al. 2019. “An Improved de Novo Assembly and Annotation of the Tomato Reference Genome Using Single- Molecule Sequencing, Hi-C Proximity Ligation and Optical Maps.” Preprint, bioRxiv, September 14. 10.1101/767764.

43. Hull-Sanders, Helen M., Micky D. Eubanks, and David E. Carr. 2005. “Inbreeding Depression and Selfing Rate of Ipomoea Hederacea Var. Integriuscula (Convolvulaceae).” American Journal of Botany 92 (11): 1871–77. 10.3732/ajb.92.11.1871.

44. Ilegems, Michael, Véronique Douet, Marlyse Meylan-Bettex, et al. 2010. “Interplay of Auxin, KANADI and Class III HD-ZIP Transcription Factors in Vascular Tissue Formation.” Development 137 (6): 975–84. 10.1242/dev.047662.

45. Iwasaki, Mayumi, and Eiji Nitasaka. 2006. “The FEATHERED Gene Is Required for Polarity Establishment in Lateral Organs Especially Flowers of the Japanese Morning Glory (I Pomoea Nil).” Plant Molecular Biology 62 (6): 913–25. 10.1007/s11103-006-9066-2.

46. Izhaki, Anat, and John L. Bowman. 2007. “KANADI and Class III HD-Zip Gene Families Regulate Embryo Patterning and Modulate Auxin Flow during Embryogenesis in Arabidopsis.” The Plant Cell 19 (2): 495–508. 10.1105/tpc.106.047472.

47. Joo, Jaehyun, and Blanca Himes. 2023. Snpsettest: A Set-Based Association Test Using GWAS Summary Statistics. https://CRAN.R-project.org/package=snpsettest.

48. Kalve, Shweta, Dirk De Vos, and Gerrit T. S. Beemster. 2014. “Leaf Development: A Cellular Perspective.” Frontiers in Plant Science 5 (July). 10.3389/fpls.2014.00362.

49. Kerstetter, Randall A., Krista Bollman, R. Alexandra Taylor, Kirsten Bomblies, and R. Scott Poethig. 2001. “KANADI Regulates Organ Polarity in Arabidopsis.” Nature 411 (6838): 706–9. 10.1038/35079629.

50. Kosentka, Pawel Z., Alexander Overholt, Richard Maradiaga, Omar Mitoubsi, and Elena D. Shpak. 2019. “EPFL Signals in the Boundary Region of the SAM Restrict Its Size and Promote Leaf Initiation.” Plant Physiology 179 (1): 265–79. 10.1104/pp.18.00714.

51. Krueger, Felix, Frankie James, Phil Ewels, et al. 2023. FelixKrueger/TrimGalore: V0.6.10 - Add Default Decompression Path. V. 0.6.10. Zenodo, released February. 10.5281/zenodo.7598955.

52. Li, Si-Bei, Zong-Zhou Xie, Chun-Gen Hu, and Jin-Zhi Zhang. 2016. “A Review of Auxin Response Factors (ARFs) in Plants.” Frontiers in Plant Science 7 (February). 10.3389/fpls.2016.00047.

53. Lincoln, C., J. Long, J. Yamaguchi, K. Serikawa, and S. Hake. 1994. “A Knotted1-like Homeobox Gene in Arabidopsis Is Expressed in the Vegetative Meristem and Dramatically Alters Leaf Morphology When Overexpressed in Transgenic Plants.” The Plant Cell 6 (12): 1859–76. 10.1105/tpc.6.12.1859.

54. Love, Jill M., and Kathleen G. Ferris. 2024. “Local Adaptation to an Altitudinal Gradient: The Interplay between Mean Phenotypic Trait Variation and Phenotypic Plasticity in *Mimulus Laciniatus*.” Perspectives in Plant Ecology, Evolution and Systematics 63 (June): 125795. 10.1016/j.ppees.2024.125795.

55. Magwene, Paul M., John H. Willis, and John K. Kelly. 2011. “The Statistics of Bulk Segregant Analysis Using Next Generation Sequencing.” PLoS Computational Biology 7 (11): e1002255. 10.1371/journal.pcbi.1002255.

56. Mansfeld, Ben N., and Rebecca Grumet. 2018. “QTLseqr: An R Package for Bulk Segregant Analysis with Next-Generation Sequencing.” The Plant Genome 11 (2): 180006. 10.3835/plantgenome2018.01.0006.

57. Marsch-Martínez, Nayelli, Víctor M. Zúñiga-Mayo, Humberto Herrera-Ubaldo, et al. 2014. “The NTT Transcription Factor Promotes Replum Development in Arabidopsis Fruits.” The Plant Journal 80 (1): 69–81. 10.1111/tpj.12617.

58. Martin, Marcel. 2011. “Cutadapt Removes Adapter Sequences from High-Throughput Sequencing Reads.” EMBnet.Journal 17 (1): 1. 10.14806/ej.17.1.200.

59. Mohr, Charles. 1901. Plant Life of Alabama: An Account of the Distribution, Modes of Association, and Adaptations of the Flora of Alabama, Together with a Systematic Catalogue of the Plants Growing in the State. Brown printing co. https://catalog.hathitrust.org/Record/102814711.

60. Nagy, Eric S. 1997. “SELECTION FOR NATIVE CHARACTERS IN HYBRIDS BETWEEN TWO LOCALLY ADAPTED PLANT SUBSPECIES.” Evolution 51 (5): 1469–80. 10.1111/j.1558-5646.1997.tb01470.x.

61. Nicotra, Adrienne B., Andrea Leigh, C. Kevin Boyce, et al. 2011. “The Evolution and Functional Significance of Leaf Shape in the Angiosperms.” Functional Plant Biology 38 (7): 535. 10.1071/FP11057.

62. Nikovics, Krisztina, Thomas Blein, Alexis Peaucelle, et al. 2006. “The Balance between the MIR164A and CUC2 Genes Controls Leaf Margin Serration in Arabidopsis.” The Plant Cell 18 (11): 2929–45. 10.1105/tpc.106.045617.

63. Okushima, Yoko, Paul J. Overvoorde, Kazunari Arima, et al. 2005. “Functional Genomic Analysis of the AUXIN RESPONSE FACTOR Gene Family Members in Arabidopsis Thaliana: Unique and Overlapping Functions of ARF7 and ARF19.” The Plant Cell 17 (2): 444–63. 10.1105/tpc.104.028316.

64. Ori, Naomi, Aya Refael Cohen, Adi Etzioni, et al. 2007. “Regulation of LANCEOLATE by miR319 Is Required for Compound-Leaf Development in Tomato.” Nature Genetics 39 (6): 787–91. 10.1038/ng2036.

65. Ostria-Gallardo, Enrique, Aashish Ranjan, Daniel H. Chitwood, et al. 2016. “Transcriptomic Analysis Suggests a Key Role for SQUAMOSA PROMOTER BINDING PROTEIN LIKE, NAC and YUCCA Genes in the Heteroblastic Development of the Temperate Rainforest Tree Gevuina Avellana (Proteaceae).” New Phytologist 210 (2): 694–708. 10.1111/nph.13776.

66. Peake, Amanda L., Emily Glasgow, Claire Abbasi, et al. 2026. “A Newly Arisen Indel Governs a Leaf Shape Polymorphism in the Ivy Leaf Morning Glory (*Ipomoea Hederacea*).” bioRxiv, January 1, 2026.06.04.730136. 10.64898/2026.06.04.730136.

67. Pekker, Irena, John Paul Alvarez, and Yuval Eshed. 2005. “Auxin Response Factors Mediate Arabidopsis Organ Asymmetry via Modulation of KANADI Activity.” The Plant Cell 17 (11): 2899–910. 10.1105/tpc.105.034876.

68. Petrášek, Jan, Jozef Mravec, Rodolphe Bouchard, et al. 2006. “PIN Proteins Perform a Rate- Limiting Function in Cellular Auxin Efflux.” Science 312 (5775): 914–18. 10.1126/science.1123542.

69. Pien, Stéphane, Joanna Wyrzykowska, Simon McQueen-Mason, Cheryl Smart, and Andrew Fleming. 2001. “Local Expression of Expansin Induces the Entire Process of Leaf Development and Modifies Leaf Shape.” Proceedings of the National Academy of Sciences of the United States of America 98 (20): 11812–17. 10.1073/pnas.191380498.

70. R Core Team. 2024. R: A Language and Environment for Statistical Computing. R Foundation for Statistical Computing. https://www.R-project.org/.

71. Richards, Thomas J., Daniel Ortiz-Barrientos, and Katrina McGuigan. 2019. “Natural Selection Drives Leaf Divergence in Experimental Populations of Senecio Lautus under Natural Conditions.” Ecology and Evolution 9 (12): 6959–67. 10.1002/ece3.5263.

72. Royer, Dana L., Jennifer C. McElwain, Jonathan M. Adams, and Peter Wilf. 2008. “Sensitivity of Leaf Size and Shape to Climate within Acer Rubrum and Quercus Kelloggii.” New Phytologist 179 (3): 808–17. 10.1111/j.1469-8137.2008.02496.x.

73. Sauer, Michael, and Jürgen Kleine-Vehn. 2019. “PIN-FORMED and PIN-LIKES Auxin Transport Facilitators.” Development (Cambridge, England) 146 (15): dev168088. 10.1242/dev.168088.

74. Scarpella, Enrico, Michalis Barkoulas, and Miltos Tsiantis. 2010. “Control of Leaf and Vein Development by Auxin.” Cold Spring Harbor Perspectives in Biology 2 (1): a001511. 10.1101/cshperspect.a001511.

75. Schrick, Kathrin, Bilal Ahmad, and Hieu V. Nguyen. 2023. “HD-Zip IV Transcription Factors: Drivers of Epidermal Cell Fate Integrate Metabolic Signals.” Current Opinion in Plant Biology 75 (October): 102417. 10.1016/j.pbi.2023.102417.

76. Shimada, Tomoo, Shigeo S. Sugano, and Ikuko Hara-Nishimura. 2011. “Positive and Negative Peptide Signals Control Stomatal Density.” Cellular and Molecular Life Sciences 68 (12): 2081–88. 10.1007/s00018-011-0685-7.

77. Shreve, Forrest, F. W. Besley, Frederick H. Blodgett, Mintin Asbury Chrysler, and Maryland State Weather Service. 1910. The Plant Life of Maryland. Special Publication / Maryland Weather Service ;3. Johns Hopkins Press. https://catalog.hathitrust.org/Record/001661617.

78. Singhal, Yash Kumar, Julia Anne Boyle, and John R. Stinchcombe. 2025. “Differences in Stomatal Conductance between Leaf Shape Genotypes of Ipomoea Hederacea Suggest Divergent Ecophysiological Strategies.” microPublication Biology 2025 (March): 10.17912/micropub.biology.001528. https://doi.org/10.17912/micropub.biology.001528.

79. Stevens, William Chase. 1948. Kansas Wild Flowers. Univ. of Kansas Press. https://catalog.hathitrust.org/Record/001494147.

80. Stinchcombe, John R. 2002. “Fitness Consequences of Cotyledon and Mature-Leaf Damage in sthe Ivyleaf Morning Glory.” Oecologia 131 (2): 220–26. 10.1007/s00442-002-0871-2.

81. Stinchcombe, John R., Mark D. Rausher, and Associate Editor: David E. McCauley. 2001. “Diffuse Selection on Resistance to Deer Herbivory in the Ivyleaf Morning Glory, Ipomoea Hederacea.” The American Naturalist 158 (4): 376–88. 10.1086/321990.

82. Tabata, Ryo, Masaya Ikezaki, Takahiro Fujibe, et al. 2010. “Arabidopsis AUXIN RESPONSE FACTOR6 and 8 Regulate Jasmonic Acid Biosynthesis and Floral Organ Development via Repression of Class 1 KNOX Genes.” Plant and Cell Physiology 51 (1): 164–75. 10.1093/pcp/pcp176.

83. Takagi, Hiroki, Akira Abe, Kentaro Yoshida, et al. 2013. “QTL-Seq: Rapid Mapping of Quantitative Trait Loci in Rice by Whole Genome Resequencing of DNA from Two Bulked Populations.” The Plant Journal 74 (1): 174–83. 10.1111/tpj.12105.

84. Tameshige, Toshiaki, Takeshi Tsuchida, Yuuki Matsushita, et al. 2025. “Mutual Inhibition between EPFL2 and Auxin Extends the Intervals of Periodic Leaf Morphogenesis.” Nature Communications 16 (1): 9753. 10.1038/s41467-025-65792-y.

85. Van der Auwera, Geraldine A., Mauricio O. Carneiro, Christopher Hartl, et al. 2013. “From FastQ Data to High-Confidence Variant Calls: The Genome Analysis Toolkit Best Practices Pipeline.” Current Protocols in Bioinformatics 43 (1): 11.10.1–11.10.33. 10.1002/0471250953.bi1110s43.

86. Vanhaeren, Hannes, Youn-Jeong Nam, Liesbeth De Milde, et al. 2017. “Forever Young: The Role of Ubiquitin Receptor DA1 and E3 Ligase BIG BROTHER in Controlling Leaf Growth and Development.” Plant Physiology 173 (2): 1269–82. 10.1104/pp.16.01410.

87. Vasimuddin, Md., Sanchit Misra, Heng Li, and Srinivas Aluru. 2019. “Efficient Architecture- Aware Acceleration of BWA-MEM for Multicore Systems.” 2019 IEEE International Parallel and Distributed Processing Symposium (IPDPS), May, 314–24. 10.1109/IPDPS.2019.00041.

88. Vogel, Steven. 1970. “Convective Cooling at Low Airspeeds and the Shapes of Broad Leaves.” Journal of Experimental Botany 21 (1): 91–101. 10.1093/jxb/21.1.91.

89. Wen, Lichao, Tao Liu, Zhichao Deng, et al. 2022. “Characterization of NAC Transcription Factor NtNAC028 as a Regulator of Leaf Senescence and Stress Responses.” Frontiers in Plant Science 13 (August). 10.3389/fpls.2022.941026.

90. Wu, Lingjie, Zhendong Tian, and Junhong Zhang. 2018. “Functional Dissection of Auxin Response Factors in Regulating Tomato Leaf Shape Development.” Frontiers in Plant Science 9 (July). 10.3389/fpls.2018.00957.

91. Wyatt, Robert, and Janis Antonovics. 1981. “Butterflyweed Re-Revisited: Spatial and Temporal Patterns of Leaf Shape Variation in Asclepias Tuberosa.” Evolution 35 (3): 529–42. 10.2307/2408200.

92. Xiong, Haiyan, Haidong He, Yu Chang, et al. 2025. “Multiple Roles of NAC Transcription Factors in Plant Development and Stress Responses.” Journal of Integrative Plant Biology 67 (3): 510–38. 10.1111/jipb.13854.

93. Xiong, Yuanyuan, Binbin Wu, Fei Du, et al. 2021. “A Crosstalk between Auxin and Brassinosteroid Regulates Leaf Shape by Modulating Growth Anisotropy.” Molecular Plant 14 (6): 949–62. 10.1016/j.molp.2021.03.011.

94. Zhu, Zhangsheng, Binmei Sun, Xiaoxia Xu, et al. 2016. “Overexpression of AtEDT1/HDG11 in Chinese Kale (Brassica Oleracea Var. Alboglabra) Enhances Drought and Osmotic Stress Tolerance.” Frontiers in Plant Science 7 (August). 10.3389/fpls.2016.01285.

